# Larvicidal evaluation of the *Origanum majorana* L. Essential Oil against the larvae of the *Aedes aegypti* mosquito

**DOI:** 10.1101/595900

**Authors:** Renata do Socorro Barbosa Chaves, Rosany Lopes Martins, Alex Bruno Lobato Rodrigues, Érica de Menezes Rabelo, Ana Luzia Ferreira Farias, Camila Mendes da Conceição Vieira Araújo, Talita Fernandes Sobral, Allan Kardec Ribeiro Galardo, Sheylla Susan Moreira da Silva de Almeida

## Abstract

This study evaluated the larvicidal activity of *O. majorana* essential oil, identified the chemical composition, evaluated the antimicrobial, cytotoxic and antioxidant potential. The larvicidal activity was evaluated against larvae of the third stage of *Aedes aegypti*, whereas the chemical composition was identified by gas chromatography coupled to mass spectrometer, the antimicrobial activity was carried out against the bacteria *Pseudomonas aeruginosa, Escherichia coli* and *Staphylococcus auereus*, the antioxidant activity was evaluated from of 2.2-diphenyl-1-picryl-hydrazila sequestration and *Artemia salina* cytotoxicity. Regarding to the results, the larvicidal activity showed that *O. majorana* essential oil caused high mortality in *A. aegypti* larvae. In the chromatographic analysis, the main component found in *O. majorana* essential oil was pulegone (57.05%), followed by the other components verbenone (16.92%), trans-p-menthan-2-one (8.57%), isomenthone (5.58%), piperitone (2.83%), 3-octanol (2.35%) and isopulegol (1.47%). The antimicrobial activity showed that *E. coli* and *P. aeruginosa* bacteria were more sensitive to oil than *S. aureus*, which was resistant at all concentrations. Essential oil did not present antioxidant activity, but it has high cytotoxic activity against *A. salina*.

## 1 INTRODUCTION

Dengue remains an important public health problem in Brazil, even after the introduction and recent dissemination of the Zika and chikungunya [1,2] viruses. The disease presents a great epidemic potential, affecting all regions of Brazil [3,4].

The *A. aegypti* mosquito (Linnaeus, 1762) is a vector of viruses that cause diseases known as dengue, chikungunya, and zika [5]. It has holometabolic development, with egg, larva, pupa and adult phases. Because it is a mosquito highly adapted to the urban environment, its most common breeding sites are artificial containers that accumulate water, such as bottles, tires, cans, and pots [6].

Among the control policies adopted in Brazil, the mechanical control is carried out by Agents to Combat Endemics (ACE), with the participation of the population, aiming at the protection, destruction or adequate allocation of potent breeding sites. The intensive collaboration of the population is crucial to hinder the proliferation and installation of the mosquito. In addition, it reinforces the need for adequate sanitary conditions in the cities, eliminating stocks of water that allow eggs to hatch. An important strategy is the promotion of educational actions during home visits made regularly by the health agents [7].

The spread and flow of various serotypes of the dengue virus over the years also have a significant influence on epidemics, as well as an increase in cases diagnosed for the most severe form of the disease. These factors demonstrate the importance of introducing preventive measures in order to reduce dengue rates [8].

*Origanum majorana* L. belongs to the Lamiaceae family, and it contains several terpenoids, which are isolated from aerial parts of the *Origanum* plant and exhibit antimicrobial, antiviral and antioxidant properties, without toxic effects [9,10]. In folk medicine, *O. majorana* is used for cramp, depression, migraine and nerve headaches [9].

The antimicrobial and antioxidant properties of many spices and their essential oils have been known for a long time, but only in recent years have consumers given proper attention to the use of these substances [11]. Because many plants are toxic to mosquitoes, the mixture of essential oils may represent an efficient outlet for this problem, compared to the *A. aegypti* mosquito [12].

In the literature, there are no reports on larvicidal activity against *A. aegypti* and cytotoxicity against *A. salina*, and few studies have been reported on the antioxidant and antimicrobial effects of the essential oils of this species.

Therefore, the objective of this study was to evaluate the larvicidal activity against *A. aegypti*, to determine the chemical composition, to evaluate the antimicrobial activity against *E. coli, P. aeruginosa* and *S. aureus* bacteria, to determine the antioxidant potential through the sequestration of DPPH and cytotoxicity against *A. salina* of *O. majorana* essential oil.

## 2 MATERIALS AND METHODS

### 2.1 Plant Material

The leaves of *O. majorana* were collected in the district of Fazendinha (00 “36’955” S and 51 “11’03’77” W) in the Municipality of Macapá, Amapá. Five samples of the plant species were deposited at the Amapaense Herbarium (HAMAB) of the Institute of Scientific Research and Technology of Amapá (IEPA).

### 2.1 Obtaining Essential Oil

The essential oil (EO) was obtained by the hydrodistillation process using the Clevenger type apparatus, 131 g of *O. majorana* dried leaves were dried at 45 °C for a period of 2 h [13]. The EO was kept under refrigeration (4°C).

### 2.3 Identification of the Chemical Composition by Gas Chromatography Coupled to Mass Spectrometer (GC-MS)

The EO analysis was performed by Gas Chromatography coupled to the Mass Spectrometer (GC-MS) of the Museu Paraense Emílio Goeldi. The Shimadzu equipment, CGEM-SHIMADZU QP 5000 was used. A fused silica capillary column (OPTIMA®-5-0.25 μm) was used. It has 30 m of length and 0.25 mm of internal diameter and nitrogen as carrier gas. The operating conditions of the gas chromatograph were: internal column pressure 67.5 kPa, division ratio 1:20, gas flow at column 1.2 mL/min (210 °C), injector temperature 260 °C, temperature detector or interface (GC-MS) of 280 °C. The initial column temperature was 50°C, followed by an increase from 6 °C/min to 260 °C kept constant for 30 min. The mass spectrometer was programmed to perform readings at intervals of 29-400 Da, 0.5 s with ionization energy of 70 eV.

The identification of the chemical compounds present in the EO was made from the comparisons of the Indices of Retention (IR) and Kovats (IK) of the homologous series of n-alkanes (C_8_-C_26_) and the literature [14]. Identification was also made by combining the spectra obtained by the analysis performed on the Labsolutions GC-MS version 2.50 SU1e software equipment of the mass spectra of the NIST05 and WILEY’S libraries.

### 2.4 Larvicidal Activity Against *A. aegypti*

The *A. aegypti* larvae used in the bioassay came from the colony kept in the Insectary of the Scientific and Technological Research Institute (IEPA). The methodology used followed the Who [15] standard with adaptations.

The procedure started with the separation of 18 beakers of 50 mL and in each Becker, there were added 25 larvae of the third instar of *A. aegypti*. Then they were reserved in a room with conditions of ambient temperature between 25 to 30 °C and photoperiod of 12 h.

Preparation of the samples started after 24 h. The stock solution was prepared with 4.5 mL of 5% Tween 80, 85.5 mL of distilled water and 0.09 g of the EO sample of *O. majorana*. The positive control was prepared with 17.5 ml of Tween 80 dissolved in 350 ml of distilled water.

In the bioassay, 18 beakers of 100 mL of glass were organized into six groups. The mother solution was distributed as follows: in group I, it was added 10 mL, in group II it was added 8 mL, in the group III it was added 6 mL, in group IV it was added 4 mL and in group V it was added 2 mL. In group VI, 80 mL of positive control solution was added.

Then, about 80% of distilled water were added to each beaker from the total volume and plus 25 *A. aegypti* larvae. Group I was 100 μg.mL^−1^, group II was 80 μg.mL^−1^, group III was 60 μg.mL^−1^, group IV was 40 μg.mL ^−1^ group V had a total of 20 μg.mL^−1^ of test solution. After 24 and 48 h, the number of dead larvae was counted, it is considered as dead all those that were unable to reach the surface.

#### 2.4.1 Larvicidal Statistical Analysis

The experiment was carried out in triplicate. The larval mortality efficiency data were calculated in percentages using the Abbott formula and later tabulated in Microsoft Excel (Version 2013 for Windows). LC_50_ (lethal concentration causing 50% mortality in the population) was analyzed by the IBM SPSS® program [version 21.0; SPSS Inc., Chicago, IL, USA]. The results were shown in the table. Differences that presented probability levels p≤0.001 were considered statistically significant.

### 2.5 Antimicrobial Activity

#### 2.5.1 Microorganisms

The antimicrobial EO test obtained from *O. majorana* leaves was tested in vitro against two gram-negative bacteria (*P. aeruginosa* ATCC 25922 and *E. coli* ATCC 8789) and a gram-positive bacterium (*S. aureus* ATCC 25922).

For each microorganism, the stock culture was stored in BHI medium (Brain Heart Infusion) with 20% glycerol and stored at −80 °C. An aliquot of 50 μL of this culture was inoculated into 5 mL of sterile BHI broth medium and incubated for 24 h at 37 °C.

#### 2.5.2 Determination of Minimum Inhibitory Concentration (MIC) and Minimum Bactericidal Concentration (MBC)

The MIC and MBC were determined using the microplate dilution technique (96 wells) according to the protocol established by Icls [16], with adaptations.

Bacteria were initially reactivated from the stock cultures, kept in BHI broth, for 18 h at 37 °C. Subsequently, bacterial growth was prepared in 0.9% saline inoculum for each microorganism, adjusted to the McFarland 0.5 scale, then diluted in BHI and tested at 2 × 10^6^ UFC.mL^−1^ concentration.

In determining the MIC, the EO was diluted in Dimethylsulfoxide (2% DMSO). Each well of the plate was initially filled with 0.1 mL of 0.9% NaCl, except for the first column, which was filled with 0.2 mL of the EO at the concentration of 2000 μg.mL^−1^. Subsequently, base two serial dilutions were performed in the ratio of 1:2 to 1:128 dilution in a final volume of 0.1 mL. After this process, 0.1 mL of cells (2 × 106 CFU.mL^−1^) added in each well related to the second preceding item, resulting in a final volume of 0.2 mL. Control of culture medium, control of EO, and negative control (DMSO 2%) were performed. And for the positive control, amoxicillin (50 μg.mL^−1^) was used. After incubation of the microplates in an incubator at 37°C for 24 hours, the plates were read in ELISA reader (DO630nm).

The determination of MBC was performed based on the results obtained in the MIC test. Microplate wells were replicated in Müller-Hinton agar and incubated at 37 °C for 24 h. MBC was established as the lowest concentration of EO capable of completely inhibiting microbial growth.

#### 2.5.3 Statistical analysis of microbiological assays

All experiments were performed in triplicate with the respective results categorized in Microsoft Excel (Version 2013 for Windows) and later analyzed in GraphPad Prism software (Version 6.0 for Windows, San Diego California USA). Significant differences between the groups were verified using the One-way ANOVA test with Bonferroni post-test. The data were considered statistically significant when p <0.001.

### 2.6 Antioxidant Activity

The antioxidant quantitative test was based on the methodology recommended by Sousa et al. [17], Lopes-Lutz et al. [18] and Andrade et al. [19] by the use of 2.2-diphenyl-1-picryl-hydrazila (DPPH) with adaptations.

A methanolic solution of DPPH (stock solution) was prepared at the concentration of 40 μg.mL^−1^, which was kept under the light. The EOs were diluted in methanol at concentrations 5, 2.5, 1.0, 0.75, 0.50 and 0.25 μg.mL^−1^. For the evaluation of the test, 0.3 mL of the oil solution was added to a test tube, followed by the addition of 2.7 mL of the DPPH solution. White was prepared from a mixture with 2.7 mL of methanol and 0.3 mL of the methanol solution of each EO concentration as measured. After 30 min. the readings were performed on a spectrophotometer (Biospectro SP-22) at a wavelength of 517 nm. The test was performed in triplicate and the calculation of the percentage of antioxidant activity (% AA) was calculated with the following equation:

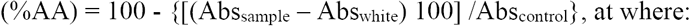

%AA - Percentage of antioxidant activity

Abs_sample_ – Sample Absorbance

Abs_white_-White Absorbance Abs_control_-Control Absorbance

### 2.7 Cytotoxic Activity Front *Artemia Salina* Leach

The *A. salina* cytotoxicity assay was based on the technique of Araújo et al. [20] and Lôbo et al. [21] with adaptations. An aqueous solution of artificial sea salt was prepared (35 gL^−1^) at pH 9.0 for incubation of 45 mg of *A. salina* eggs, which were placed in the dark for 24 h for the larvae to hatch (nauplii), then the nauplii were exposed to artificial light in 24 h, period to reach the stage methanuplii. The stock solution was prepared to contain 0.06 g of EO, 1.5 mL of Tween 80 and 28.5 mL of saline to facilitate solubilization of the same. The test tubes were marked up to 5 mL.

The methanauplia were selected and divided into 7 groups of 10 subjects in each test tube. Each group received aliquots of the stock solution (2500, 1875, 1250, 625, 250, and 125 μL), which were then filled to a volume of 5 mL with the sea salt solution to produce final solutions with the following concentrations: 1000, 750, 500, 250, 100 and 50 μg.mL^−1^. The tests were performed in triplicates. For the test control, saline solution was used. After 24 h, the number of dead was counted. The lethal concentration causing 50% mortality in the population (LC_50_) was determined by Probit analysis using SPSS Software (version 20.0; SPSS Inc., Chicago, IL, USA).

## 3 RESULTS

### 3.1 Identification of Chemical Compounds by GC-MS of *O. majorana* Essential Oil

The chemical composition was determined by GC-MS, where the chromatogram can be observed in Fig 1. On the chemical composition of the EO of *O. majorana* (Table 1), 95.8% are oxygenated monoterpenes and only 1.38% are monoterpene hydrocarbons. The major component of EO is pulegone (57.05%), followed by other components verbenone (16.92%), trans-menthone (8.57%), cis-menthone (5.58%), piperitone (2.83%), 3-octanol %) and isopulegol (1.47%).

**Fig. 1.**
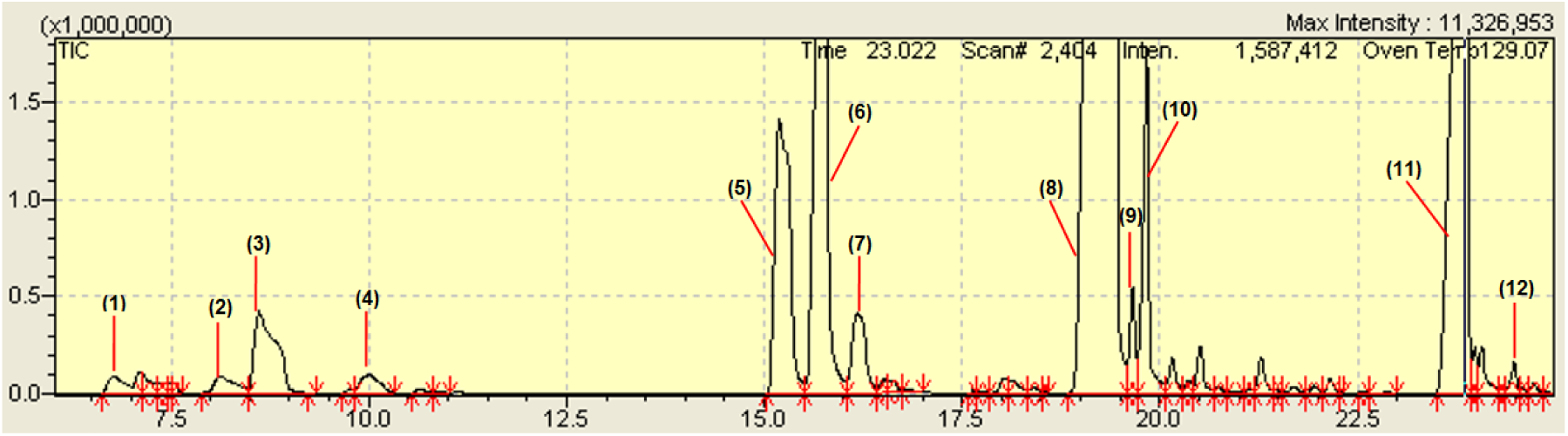
Obtaining Gas Chromatography of *O. majorana* essential oil. Notes: Gas Trap: Helium (He); initial temperature 60 ° C; initial time 1.0 min; the column temperature increased 3 ° C / min. at 240 ° C, maintained at this temperature for 30 min.

**Tabela1.** Chemical composition of *O. majorana* essential oil.

| Nº | IR | IK | Compounds | Relative Percentage (%) | Identification * |
| --- | --- | --- | --- | --- | --- |
| 1 | 6.776 | 939 | α-pinene | 0.39 | MS, IK |
| 2 | 8.127 | 979 | β-pinene | 0.50 | MS IK |
| 3 | 8.617 | 991 | 3-octanol | 2.35 | MS, IK |
| 4 | 9.995 | 1029 | Limonene | 0.49 | MS, IK |
| 5 | 15.185 | 1162 | iso-menthone | 5.58 | MS, IK |
| 6 | 15.688 | 1199 | trans-*p*-menthan-2-one | 8.57 | MS, IK |
| 7 | 16.174 | 1149 | Isopulegol | 1.47 | MS, KI |
| **8** | **19.448** | **1237** | **Pulegone** | **57.05** | **MS, IK** |
| 9 | 19.655 | 1165 | Lavandulol | 0.77 | MS, IK |
| 10 | 19.883 | 1252 | Piperitone | 2.83 | MS, IK |
| 11 | 23.873 | 1205 | Verbenone | 16.92 | MS, IK |
| 12 | 24.482 | 1161 | nonen-1-al-(2E) | 0.26 | MS, IK |
|  |  |  | monoterpenes hydrocarbon | 95.8 |  |
|  |  |  | monoterpenes oxide | 1.38 |  |
|  |  |  | Total | 97.18 |  |

Notes: tR: retention time. The identification path of the compounds, the identification of the compounds was performed by the mass spectrum (MS) of the Labsolutions CG-EM software version 2.50 SU1 (NIST05 and WILEY'S libraries of the 9th edition mass spectrum) and Kovats Index (KI) (ADAMS, 2017).

### 3.2 Larvicidal Activity

The percentage of dead *A. aegypti* larvae is shown in Table 2, at different EO concentrations of *O. majorana* in the 24-48 h exposure period. There was no mortality in the control group. Through the Probit test, LC_50_ = 56.008 μg.mL^−1^, determination coefficient (R^2^) = 0.917 and quantitative evaluation (*X*_*2*_) = 0.844 in 24 h. After 48 h at LC_50_ = 15.696 μg.mL^−1^, *X*^*2*^ = 0.572 and R^2^ = 0.835.

**Tabela 2.** Porcentagem de larvas mortas (%) de *A. aegypti* produzida por diferentes concentrações do óleo essencial de *O. majorana* em 24-48 h.

| Concentrações ($\mu\text{g.mL}^{-1}$) | 24 h | 48 h |
| --- | --- | --- |
| Controle | 0 | 0 |
| 20 | 16* | 57.33* |
| 40 | 40* | 76* |
| 60 | 40* | 77.3*3 |
| 80 | 65.32* | 82.66* |
| 100 | 78.62* | 94.66* |

Fonte: dados do estudo; \* letras deferentes indicam uma diferença significativa $p < 0.0001$ .

### 3.3 Antimicrobial Activity

The minimum inhibitory concentration (MIC) and minimum bactericidal concentration (MBC) that were identified for O. majorana EO can be verified in Fig. 2. The results show that gram-negative bacteria were more sensitive to EO presenting MIC = 31.25 μg.mL^−1^ compared to the negative control. The MBC for *E. coli* was at the concentration of 500 μg.mL^−1^ and for *P. aeruginona* was at the concentration of 1000 μg.mL^−1^ in relation to the negative control (amoxicillin). While the *S. aureus* bacterium did not present MIC and neither MBC.

**Fig. 2.**
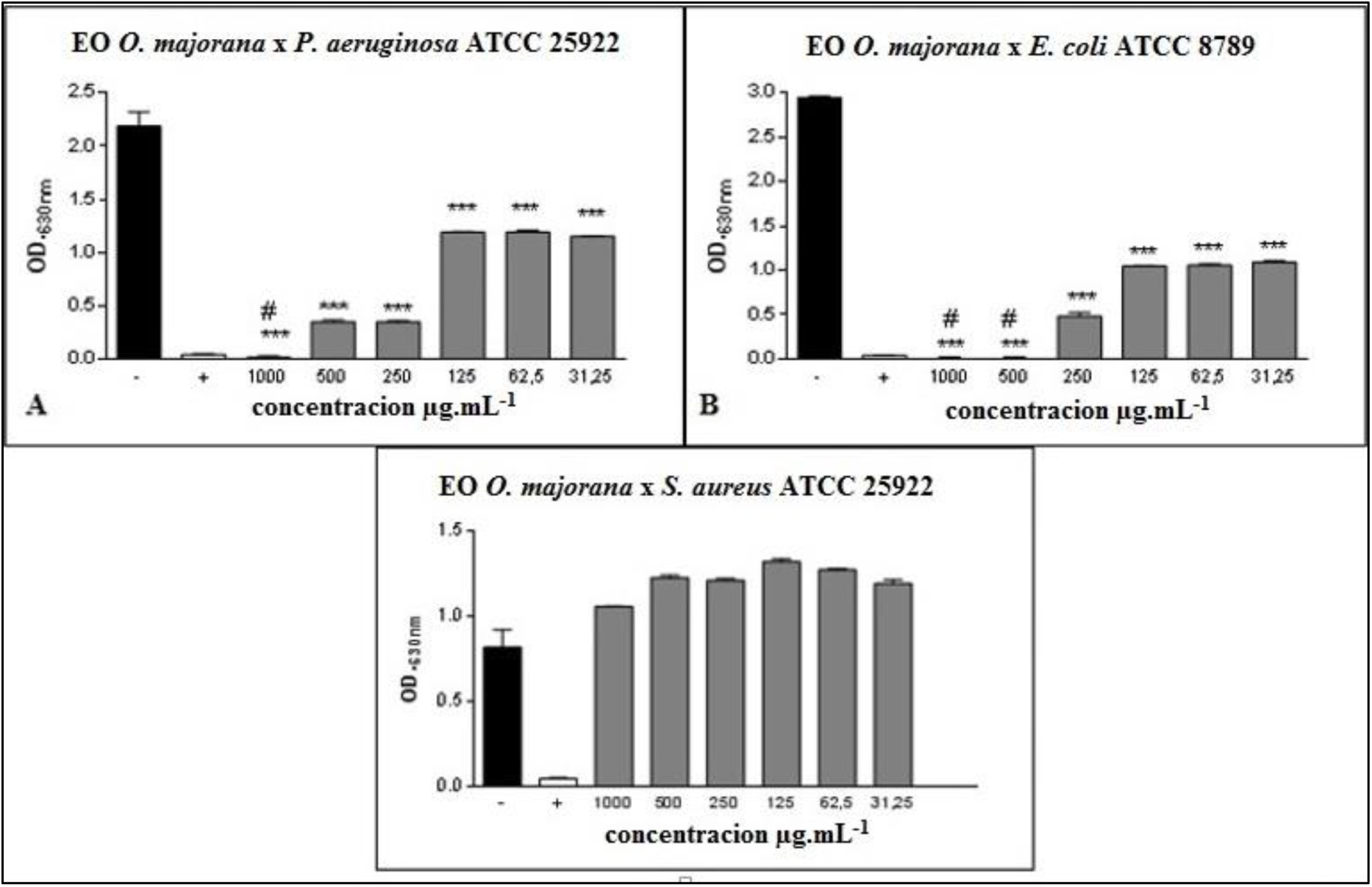
Concentração Inibitória Mínima (CIM) e Concentração Bactericida Mínima (CBM) do óleo essencial de *Origanum majorana*. Fonte: dados do estudo, substância teste 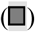, ICC com 2% DMSO (▪) e Amoxilina (□); *P<0.001 estatisticamente significante em relação ao controle negativo; # p<0.001 estatisticamente significante em relação ao controle positivo; e (D) Método de difusão em disco do OE de *O. majorana*.

### 3.4 Antioxidant Activity

Table 3 shows the percentage of antioxidant activity of *O. majorana* EO. For the EO concentrations IC_50_ = 16.83 μg.mL^−1^ was compared with ascorbic acid (vitamin C) which showed IC_50_ = 16.71 μg.mL^−1^ as shown in Table 4. The absorbance of EO was Y = 0.0196x + 17.0078 and the coefficient (R^2^) = 0.9600.

**Tabela 3.** Média e desvio padrão do percentual da atividade antioxidante do óleo essencial de *O. majorana* em diferentes concentrações.

| Concentrações ($\mu\text{g.mL}^{-1}$) | % AA |
| --- | --- |
| 7.81 | $16.95 \pm 0.20^a$ |
| 15.62 | $17.15 \pm 0.17^b$ |
| 31.25 | $17.48 \pm 0.37^b$ |
| 62.5 | $18.98 \pm 0.56^c$ |
| 125 | $19.28 \pm 0.75^d$ |
| 250 | $21.85 \pm 2.55^e$ |

Fonte: dados do estudo; letras diferentes indicam que houve diferença significativa teste de Tukey ( $p \leq 0,05$ ).

**Tablea 4.** Média e desvio padrão do percentual de atividade antioxidante do ácido ascórbico (vitamina C) em diferentes concentrações.

| Concentrações ($\mu\text{g.mL}^{-1}$) | % AA |
| --- | --- |
| 7.81 | $18.57 \pm 0.52^a$ |
| 15.62 | $30 \pm 0.10^b$ |
| 31.25 | $99.93 \pm 0.02^c$ |
| 62.5 | $99.99 \pm 0.0^d$ |
| 125 | $99.99 \pm 0.0^d$ |
| 250 | $99.99 \pm 0.0^d$ |

Fonte: dados do estudo; os valores correspondem à média e à duração padrão das triplicatas; letras diferentes indicam que houve diferença significativa ( $p \leq 0,05$ ).

### 3.5 Cytotoxic activity

Table 5 shows the percentage of cytotoxic activity of *O. majorana* EO and the mean mortality readings after the 24 h period are expressed. The oil concentrations presented LC_50_ of 172.6 μg.mL^−1^, the coefficient of determination (R^2^) of 0.883 and *X*^*2*^ of 1.915.

**Tabela 5.**
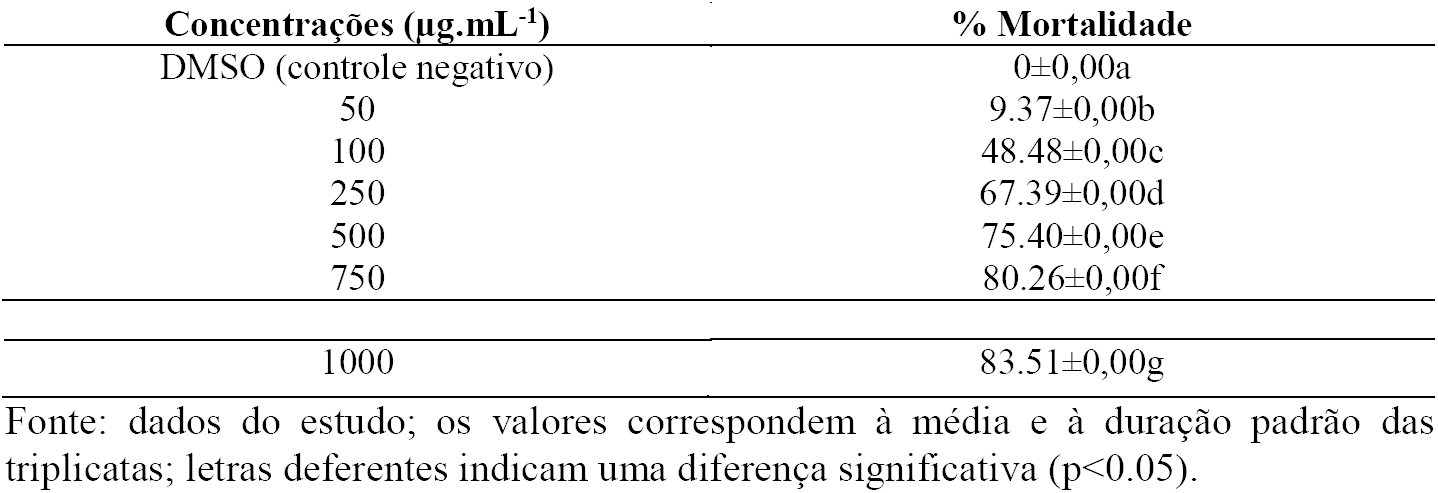
Média e desvio padrão do percentual de atividade citotóxica do óleo essencial de *O. majorana* em diferentes concentrações.

| Concentrações ($\mu\text{g.mL}^{-1}$) | % Mortalidade |
| --- | --- |
| DMSO (controle negativo) | $0 \pm 0,00a$ |
| 50 | $9.37 \pm 0,00b$ |
| 100 | $48.48 \pm 0,00c$ |
| 250 | $67.39 \pm 0,00d$ |
| 500 | $75.40 \pm 0,00e$ |
| 750 | $80.26 \pm 0,00f$ |
| 1000 | 83.51±0,00g |

Fonte: dados do estudo; os valores correspondem à média e à duração padrão das triplicatas; letras diferentes indicam uma diferença significativa (p<0.05).

## 4. DISCUSSION

This result corroborates with other studies that have shown that environmental factors may affect certain chemical compounds, while in others they have no influence on their production [22,23].

Lima et al. [24], reports that piperitone has three organic functions in its chemical structure and it can be used for the synthesis of other compounds. Piperitone is derived from the metabolic pathway for the formation of piperitenone oxide, in which cis-pulegone is also, derived [25].

Macêdo et al. [26] observe that the variations of the active components of the plant are important parameters to correlate the activities, such as antibacterial and insecticide. In addition, a number of biotic factors such as plant/ microorganism Stoppacher et al. [27], plants/ insects Kessler and Baldwin [28] plant interactions, age and stage of development. As well as abiotic factors such as luminosity Takshak and Agrawal [29], temperature, precipitation, nutrition, time and harvest time Bitu et al. [30], they may present correlations with each other, acting together, and they may exert a joint influence on chemical variability and yield of essential oil [26].

The results of the larvicidal activity of this study show that *O. majorana* EO is active against *A. aegypti* larvae. A fact that Komalamisra et al. [31], Magalhães et al. [32] and Dias et al. [33], classified with the values of the minimum lethal concentration that eliminates 50% of the population (LC_50_) as a criterion for the activity. Because if LC_50_<50 μg.mL^−1^, the product is considered very active, if 50 <LC_50_ <100 μg.mL^−1^ the product is considered active, and when LC_50_> 750 μg.mL^−1^ the product is considered inactive. There were no reports of studies on the larvicidal activity of *O. majorana* essential oil against *A. aegypti* larvae.

According to Cantrell et al. [34], larvicidal compounds act by absorption through the cuticle, via the respiratory tract, and/or enter by ingestion via the gastrointestinal tract. Once inside the larva, the substances may reach the site of action or may cause systemic effects by diffusion in different tissues [35].

Studies on the insecticidal effect of *Mentha* spp. reported that menthol, mentona, pulegone and carvone help to clarify the mechanisms of action on insects [36]. Previous studies indicate that limonene, camphene, and verbenone have insecticidal insect activity [37].

Some EOs are known to cause dissuasive or anti-eating behavior in insects suggesting a neurotoxic action Satyan et al. [38], while some act as growth-regulating insects through analogous effects or antagonistic endogenous hormones. In the present study, it was found that even short-term exposure of larvae to lethal doses can dramatically increase their mortality over time and thereby reduce the total number of viable adults, leading to a possible reduction in total populations [39].

In relation to the microbiological activity, it was possible to verify that gram-negative bacteria were more sensitive to *O. majorana* EO than gram-positive bacteria.

According to Rosato et al. [40], the antibacterial activity in gram-negative bacteria occurs due to the high percentage of oxygenated monoterpenes present in the EO and consequently the synergism between these components. On the other hand, bacteria can also respond to adverse conditions in a transient way, through so-called stress tolerance responses. Bacterial stress tolerance responses include structural and physiological modifications in the cell, and complex genetic regulatory machines mediate them [41].

In the study by Duru et al. [42] pulegone showed high antimicrobial activity, particularly against Candida, albicans and Salmonella typhimurium. Pulegone is classified as a monoterpene, in the same way as carvone. It can be obtained from a variety of plants [43,44]. Menthone is a common volatile compound in Lamiaceae, which may also be active against a large number of bacteria, such as E. coli and Enterococcus faecalis [45,46]. Some studies have argued that monoterpenes can cross cell membranes and interact with intracellular sites critical for antibacterial activity [47].

However, reports of non-adaptation or cross-adaptation of bacteria to sublethal concentrations of major constituents of essential oils have also been reported [48]. Cross-resistance can occur when different antimicrobial agents attack the same target in the cell, reach common route of access to the respective targets or initiate a common pathway for cell death, ie, the resistance mechanism is the same for more than one antibacterial agent [49].

Many antioxidants derived from natural products demonstrate neuroprotective activity in vitro and/or in vivo models such as flavonoid phenolic compounds [50].

The percentage of antioxidant activity of the essential oil showed a high IC_50_ =μg.mL^−1^, whereas ascorbic acid presented 16.71 μg.mL^−1^[51]. According to Rodrigues [52] the higher the consumption of DPPH for a smaller sample will be its IC_50_ and the greater its antioxidant capacity.

According to Beatović et al. [53], the antioxidant capacity of EO is related to its main compounds. However, this study did not present antioxidant activity. The importance concerning the performance of antioxidants depends on the factors types of free radicals formed; where and how these radicals are generated; analysis and methods for identifying damage, and ideal doses for protection [54].

*A. salina* is a microcrustacean used in fish feed, and it is widely used in toxicological studies because of the low cost and easy cultivation. Several studies have attempted to correlate toxicity on *A. salina* with antifungal, virucidal, antimicrobial, trypanosomicidal and parasiticidal activities. Lethality assays are performed in toxicological tests and the median lethal concentration (LC_50_), which indicates death in half of a sample, can be obtained [55].

According to Nguta et al. [56], both organic extracts and aqueous extracts with LC_50_ values of less than 100 μg.mL^−1^ show high toxicity, LC_50_ between 100 and 500 μg.mL^−1^ exhibited moderate toxicity, LC_50_ between 500 and 1000 μg.mL^−1^ presented low toxicity and LC_50_ above 1000 μg.mL^−1^ are considered to be non-toxic (non-toxic).

The lethal concentration of mortality against the *A. salina* larvae of this assay showed moderate cytotoxic activity. In order to evaluate the cytotoxicity of a given sample, it is possible to elucidate the cytotoxic effect of the cytotoxic mechanism and the mechanism of action of different compounds during their interaction with the tissues [57].

## CONCLUSION

The results of the present study demonstrated that OE obtained from dry leaves of *O. majorana* showed good larvicidal activity against *A. aegypti* larvae with mortality from the concentration of 20 μg.mL^−1^ in 48 h. In relation to the chemical analysis, it presented a mixture of monoterpenes, with the major constituent being pulegone (57.05%), followed by the other constituents verbenone (16.92%), trans-menthone (8.57%), cis-menthone), piperitone (2.83%), 3-octanol (2.35%) and isopulegol (1.47%). The oil showed satisfactory antimicrobial activity against *P. aeruginosa* and *E. coli* bacteria. In addition, despite the lack of antioxidant activity by the DPPH radical capture method, the oil showed moderate cytotoxic activity against *A. salina*. In summary, these results provide subsidies for future EO *O. majorana* studies in order to enhance the use of organic compounds with larvicidal activity against the *A. aegypti* mosquito, as well as the importance of the study of bioactive plant products that do not pollute the environment and that do not cause harm to human health.

## Acknowledgments

To the Amapá Foundation for Research Support (FAPEAP). To the Research Program of SUS - PPSUS - Ministry of Health. To the Coordination of Improvement of Higher Education Personnel (CAPES)/Ministry of Education (MEC). To the National Council of Scientific and Technological Development - CNPQ. To the Laboratory of Microbiology (LEMA) of UNIFAP under the responsibility of Prof. Aldo Proietti Aparecido Júnior. To the Pro-rector of research and post-graduation - PROPESPG. Federal University of Amapá UNIFAP. To the Laboratory Adolpho Ducke under the responsibility of Eloise Andrade.

## References

1. paho.org [internet] World Health Organization: Neglected, tropical and vector borne disease - dengue. c2017 [cited 2018 dec 28]. Available from: http://www.paho.org/hq/index.php?option=com_topics&view=article&id=1&Itemid=40_734.

2. Bhatt S, Gething PW, Brady OL, Messina JP, Farlow AW, Moyes CL, et al. The global distribution and burden of dengue. Nature. 25 of april of 2013; 496 (7446): 504–7. PubMed PMID: 23563266; PubMed Central PMCID: PMC3651993; doi: 10.1038/nature12060

3. Gonçalves NVS, Rebêlo JMM. [Epidemiological aspects of dengue in the municipality of São Luís, Maranhão, Brazil, 1997-2002]. Cadernos de Saúde Pública. 2004; 20: 1424–1431. Doi: 10.1590/S0102-311X2004000500039. Português.

4. Barbosa IR, Araújo LF, Carlota FC, Araújo RS, Maciel IJ. [Epidemiology of dengue fever in the State of Rio Grande do Norte, Brazil, 2000 to 2009]. Epidemiol Serv Saúde. 2012; 21: 149–57. Doi:V10.5123/S1679-49742012000100015. Português.

5. Vasconcelos PFC. Doença pelo vírus Zika: um novo problema emergente nas Américas? Rev. PanAmaz. Saúde. 2015; 6: 9–10. Doi: 10.5123/S2176-62232015000200001. Português.

6. Lima-Camara, T. N. [Emerging arboviroses and new challenges for public health in Brazil]. Ver Saúde Pública. 7 of march of 2016; 50 (36): 1–7. DOI: 10.1590/S1518-8787.2016050006791.

7. Zara A. L.; Santos, S. M. dos; Oliveira-Fernandes, E. S.; Carvalho, R. G.; Coelho, G. E. [Strategies for controlling Aedes aegypti: a review]. Epidemiol Serv Saúde. 2016; 25 (2): 391–404. Doi: 10.5123/s1679-49742016000200017. Português.

8. Araújo VEM, Bezerra JMT, Frederico FA, Passos VMA, Carneiro M. [Increased burden of dengue in Brazil and federated units, 2000 and 2015: analysis of the *Global Burden of Disease Study* 2015]. Revista Brasileira de Epidemiologia. 2017; 20: 205–216. Doi: 10.1590/1980-5497201700050017. Português.

9. Mitropoulou G, Fitsiou E, Stavropoulou E, Papavassilopoulou E, Vamvakias M, Pappa A, Oreopoulou A, Kourkoutas Y. Composition, antimicrobial, antioxidant, and antiproliferative activity of *Origanum dictamnus* (dittany) essential oil. Microbial Ecology in Health and Disease. 6 of may of 2015; 26: (1-9): 26543. PMID: 25952773; PMCID: PMC4424236; Doi: 10.3402/mehd.v26.26543.

10. Bina F, Rahimi R. Sweet Marjoram: A Review of Ethnopharmacology, Phytochemistry, and Biological Activities. J Evid Based Int Med. 2016; 22 (1): 175–185. PMID: 27231340; PMCID: PMC5871212; Doi: 10.1177/2156587216650793.

11. Dussault D, Vu KD, Lacroix M. *In vitro* evaluation of antimicrobial activities of various comercial essential oils, oleoresin and pure compounds against food pathogens and application in ham. Meat Science. 2014; 96 (1): 514–520. PMID: 24012976; Doi: 10.1016/j.meatsci.2013.08.015.

12. Pereira ÁIS, Pereira AGS, Sobrinho OPL, Cantanhede EKP, Siqueira LFS. [Antimicrobial activity in the control of larvae of the mosquito *Aedes aegypti*: Homogenization of the essential oils of linalool and eugenol]. Educ. quím. 2014; 25 (4): 446–449. Doi: 10.1016/S0187-893X(14)70065-5. Português.

13. Anvisa. National Health Surveillance Agency. [Brazilian Pharmacopoeia], 5a ed.; Fiocruz: Brasília, Brasil 2010, pp. 1–545. Português.

14. Adams RP. Identification of Essential Oil Components by Gas Chromatography/Mass Spectrometry, 4.1 ed. Biology Department: Baylor Universit, 2017.

15. World Health Organization (WHO) [internet] Geneva: Guidelines for Laboratory and Field Testing of Mosquito Larvicides. 2005c – [cited 4 december 2018]. Available on: http://apps.who.int/iris/bitstream/10665/69101/1/WHO_CDS_WHOPES_GCDPP_2005.13.pdf.

16. Clinical and Laboratory Standards Institute (CLSI) [internet] Pennsylvania: Standards for Antimicrobial Susceptibility Testing supplement M100. 2018 – [cited 5 january 2019]. Available on: https://clsi.org/media/1930/m100ed28_sample.pdf.

17. Sousa CMM, Silva HR, Junior GMV, Ayres MCC, Costa CLS, Araújo DS, Cavalcante LCD, Barros EDS, Araújo PBM, Brandão MS, Chaves MH. [Total phenols and antioxidant activity of five medicinal plants]. Química Nova. 2007; 30 (2): 351–355. Doi: 10.1590/S0100-40422007000200021. Português.

18. Lopes-Lutz D, Alviano DS, Alviano CS, Kolodziejczyk PP. Screening of chemical composition, antimicrobial and antioxidant activities of Artemisia essential oils. Phytochemistry. 2008; 69 (8): 1732–8. PMID: 18417176; DOI: 10.1016/j.phytochem.2008.02.014.

19. Andrade MA, Cardoso GM, Batista RL, Mallet TCA, Machado FMS. [Essential Oils of *Cymbopogon Nardus, Cinnamomum Zeylanicum* and *Zingiber Officinale*: composition, antioxidant and antibacterial activities]. J Agron Sci. 2012; 43 (2): 399–408. Doi: 10.1590/S1806-66902012000200025. Português.

20. Araújo MGF, Cunha RW, Veneziani RCS. Preliminary phytochemical study and toxicological bioassay against larvae of *Artemia salina* Leach. of extract obtained from fruits of *Solanum lycocarpum* A. St.-Hill (Solanaceae). J Basic Appl Pharm Sci [intenet]. 01 may 2010 [cited on 12 february 2019]; 31 (2): 205–209. Available on: https://repositorio.unesp.br/handle/11449/71676.

21. Lôbo KMS, Athayde ACR, Silva AMA, Rodrigues FFG, Lôbo IS, Bezerra DAC, Costa JGM. Evaluation of the antibacterial activity and phytochemical prospection of Solanum paniculatum Lam. and Operculina hamiltonii (G. Don) D. F. Austin e Staples, from the semi-arid region of Paraíba. Braz J Med Plants. 2010; 12 (2): 227–233. Doi: 10.1590/S1516-05722010000200016.

22. Araújo TAS, Castro VTNA, Solon LGS, Silva GA, Almeida MG, Costa JGM, Amorim ELC, Albuquerque UP. Does rainfall affect the antioxidant capacity and production of phenolic compounds of an important medicinal species?. Ind Crops Prod. 15 december 2015; 76: 550–556, 2015. Doi: 10.1016/j.indcrop.2015.07.008.

23. Estell RE, Fredrickson EL, James DK. Effect of light intensity and wavelength on concentration of plant secondary metabolites in the leaves of Flourensia cernua. Biochem Syst Ecol. 2016 april; 65: 108–114. Doi: 10.1016/j.bse.2016.02.019.

24. Lima TC, Silva TK, Silva FL, Barbosa JMJ, Marques MO, Santos RL, Cavalcanti SC, Sousa DP. Atividade larvicida do óleo essencial de *Mentha × villosa* Hudson, rotundifolona e derivados. Chemosphere. 2014 jun; 104: 37–43. Doi: 10.1016/j.chemosphere.2013.10.035.

25. Souza AVV, Santos US, Carvalho JRS, Barbosa BHS Canuto KM, Rodrigues THS. Chemical Composition of Lippia schaueriana Essential Leaf Fight Mart. Collected in the Caatinga Area. Molecules. [preprint]. 2018. [posted 2018 september 27; accepted 2018 july 6; reiceved 2018 may 23]: [6p.] Available from: https://ainfo.cnptia.embrapa.br/digital/bitstream/item/183650/1/Lippia-schaueriana-molecules-23-02480-3.pdf. Doi: 10.3390/molecules23102480

26. Macêdo DG, Souza MA, Morais-Braga MFB, Coutinho HDM, Santos ATL, Cruz RP, Costa JGM, Rodrigues FFG, Quintans JRL, Almeida JRGS, Menezes IR. Effect of seasonality on chemical profile and antifungal activity of essential oil isolated from leaves Psidium salutare (Kunth) O. Berg. Peer J. 1 november 2018; 6 (1-19). PubMed: 30402343. PMCID: PMC6215697; Doi: 10.7717/peerj.5476.

27. Stoppacher N, Kluger B, Zeilinger S, Krska R, Schuhmacher R. Identification and profiling of volatile metabolites of the biocontrol fungus Trichoderma atroviride by HS-SPME-GC-MS. J Microbiol Methods. 2010; 81 (2): 187–193. Doi: 10.1016/j.mimet.2010.03.011; PMID: 20302890.

28. Kessler A, Baldwin IT. Defensive function of herbivore-induced plant volatile emissions in nature. Science. 2001; 291 (5511): 2141–4. PMID: 11251117; Doi: 10.1126/science.291.5511.2141.

29. Takshak S, Agrawal SB. The role of supplemental ultraviolet-B radiation in altering the metabolite profile, essential oil content and composition, and free radical scavenging activities of Coleus forskohlii, an indigenous medicinal plant. Environ Sci Pollut Res Inst. 2016; 23 (8): 7324–37. PMID: 26681329; Doi: 10.1007/s11356-015-5965-6.

30. Bitu V, Costa J, Rogrigues F, Colares A, Coutinho H, Botelho M, Portela A, Santana N, Menezes I. Effect of collection time on composition of essential oil of *LippiagracilisSchauer* (Verbenaceae) growing in Northeast Brazil. Journal of Essential Oil Bearing Plants. [preprint]. 2015. [posted 2016 jul 09; received 2012 aug 08; cited 2019 jan 21]: [647–653 p.] Available from: https://www.tandfonline.com/doi/abs/10.1080/0972060X.2014.935043. doi: 10.1080/0972060X.2014.935043

31. Komalamisra N, Trongtokit Y, Rongsriyam Y, Apiwathnasorn C. Screening for larvicidal activity in some Thai plants against four mosquito vector species. Southeast Asian J Trop Med Public Health. 2005; 36 (6): 1412–22. PMID: 16610643.

32. Magalhães LAM, Lima MP, Marques MOM, Facanali R, Pinto ACS, Tadei WP. Chemical composition and larvicidal activity against *Aedes aegypti* larvae of essential oils from four Guarea species. Molecules. 2010; 15 (8): 5734–5741. PMID: 20724962; PMCID: PMC6257719; DOI: 10.3390/molecules15085734.

33. Dias CN, Moraes DFC. Essential oils and their compounds as *Aedes aegypti* L. (Diptera: Culicidae) larvicides: review. Parasitol Res. 2014; 113 (2): 565–92. PMID: 24265058; Doi: 10.1007/s00436-013-3687-6.

34. Cantrell CL, Pridgeon JW, Fronczek FR, Becnel JJ. Structure activity relationship studies on derivatives of Eudesmanolides from Inula helenium as toxicants against *Aedes aegypti* larvae and adults. Chem Biodivers. 2010; 7 (7): 1681–97. PMID: 20658657; Doi: 10.1002/cbdv.201000031.

35. Souza TM, Cunha AP, Farias DF, Machado LK, Morais SM, Ricardo NMPS, Carvalho AFU. Insecticidal activity against Aedes aegypti of m-pentadecadienyl-phenol isolated from Myracrodruon urundeuva seeds. Pest Manag Sci. 2012; 68 (10): 1380–4. PMID: 22689540; Doi: 10.1002/ps.3316.

36. Kumar P, Mishra S, Malik AS. Insecticidal properties of *Mentha* species: A review. Ind Crops Prod. 2011 jul; 34: 802–817. Doi: 10.1016/j.indcrop.2011.02.019.

37. Omolo MO, Okinyo D, Ndiege IO, Lwande W, Hassanali A. Fumigant toxicity of the essential oils of some African plants against *Anopheles gambiae* sensu stricto. Phytomedicine. 2005; 12 (3):241–6. PMID: 15830848; Doi: 10.1016/j.phymed.2003.10.004.

38. Satyan RS, Malarvannan S, Eganathan P, Rajalakshmi S, Parida A. Growth inhibitory activity of fatty acid methyl esters in the whole seed oil of *Madagascar periwinkle* (Apocyanaceae) against *Helicoverpa armigera* (Lepidoptera: Noctuidae). J Econ Entomol. 2009; 102 (3): 1197–202. PMID: 19610438.

39. Chere JMC, Dar MA, Pandit RS. Evaluation of Some Essential Oils against the Larvae of House Fly, Musca domestica by Using Residual Film Method. Biotechnol Microb. 16 april 2018; 9 (1): 555752. Doi: 10.19080/AIBM.2018.09.555752

40. Rosato A, Carocci A, Catalano A, Clodoveo ML, Franchini C, Corbo F, Carbonara GG, Carrieri A, Fracchiolla G. Elucidation of the synergistic action of Mentha Piperita essential oil with common antimicrobials. PLoS One. 01 january 2018; 13 (8):e0200902. PMID: 30067803; PMCID: PMC6070247. Doi: 10.1371/journal.pone.0200902.

41. Alvarez-Ordóñez, A.; Broussolle, V.; Colin, P.; Nguyen-The, C.; Prieto, M. The adaptive response of bacterial food-borne pathogens in the environment, host and food: implications for food safety. Int J Food Microbiol. 20 november 2015; 213: 99–109. PMID: 26116419. Doi: 10.1016/j.ijfoodmicro.2015.06.004.

42. Duru ME, Ozturk M, Ugur AU, Ceylan O. The constituents of essential oil and in vitro antimicrobial activity of *Micromeria cilicica* from Turkey. J Ethnopharmacol. 94 (1): 43–48. Doi.org/10.1016/j.jep.2004.03.053.

43. Siano F, Catalfamo M, Cautela D, Servillo L, Castaldo D. Analysis of pulegone and its enanthiomeric distribution in mint-flavoured food products. Food Addit Contam. 2005; 22 (3): 197–203. PMID: 16019787. Doi: 10.1080/02652030500041581.

44. Burt S. Essential oils: their antibacterial properties and potential applications in foods-a review. Int J Food Microbiol. 01 august 2004; 94 (3): 223–253. PMID: 15246235. Doi: 10.1016/j.ijfoodmicro.2004.03.022.

45. Singh R, Shushni MAM, Belkheir A. Antibacterial and antioxidant activities of *Mentha piperita* L. Arab J Chem. May 2015; 8 (3): 322–328. Doi.org/10.1016/j.arabjc.2011.01.019.

46. Thosar N, Basak S, Bahadure RN, Rajurkar M. Antimicrobial efficacy of five essential oils against oral pathogens: an in vitro study. Eur J Dent. September 2013; 7 (1): 71–77. PMCID: PMC4054083; PMID: 24966732. doi: 10.4103/1305-7456.119078.

47. Turchi B, Mancini S, Pistelli L, Najar B, Cerri D, Fratini F. Sub-inhibitory stress with essential oil affects enterotoxins production and essential oil susceptibility in Staphylococcus aureus. Nat Prod Res. Mach 2017; 32 (6): 682–688. PMID: 28595460. Doi: 10.1080/14786419.2017.1338284.

48. Santos JMP. [Adaptation and Cross-Adaptation of Listeria Spp. Essential Oils from Condensed Plants and Acid Stress]. [dissertação]. Programa de Pós-Graduação em Plantas Medicinais, Aromáticas e Condimentares: Universidade de Lavras; 2018. Português.

49. Chapman JS. Desinfectant resistence mechanisms, cross-resistance, and coresistance. International Int Biodeterior Biodegradation. June 2003; 51 (1): 271–276. doi:https://doi.org/10.1016/S0964-8305(03)00044-110.1016/S0964-8305(03)00044-1.

50. Kelsey N. A.; Wilkins H. M.; Linseman, D. Nutraceutical Antioxidants as Novel Neuroprotective Agents. Molecules. 03 november 2010; 15 (11): 7792–814. PMID: 21060289; PMCID: PMC4697862. Doi: 10.3390/molecules15117792.

51. Godói AA, Ishikawa BR, Ark porto, Roel RA, Xavier NCP, Yano M. [Evaluation of antioxidant, antibacterial and cytotoxic activity of *Urera aurantiaca*]. Rev Bras Farm [internet]. August 2011 [cited on 22 january 2019]; 92 (3): [198-202 2011]. Available on: http://www.rbfarma.org.br/files/rbf-2011-92-3-19.pdf.

52. Rodrigues JSQ. [Infusions based on leaves of passifloras of the cerrado: phenolic compounds, antioxidant activity in vitro and sensorial profile]. [dissertação] Universidade de Brasília. 2012. Português.

53. Beatovic D, Krstic-Miloševic D, Trifunovic S, Šiljegovic J, Glamoclija J, Ristic ME. Chemical composition, antioxidant and antimicrobial activities of the essential oils of twelve cultivars of *Ocimum basilicum* L. grown in Serbia. Rec Nat Prod. [received 2012 augu 24; revised 2013 octo 22; cited 22 january 2019]: [62-75]. Available on: https://www.acgpubs.org/doc/201808071238455-RNP-EO_1208-054.pdf.

54. Singh B, Singh JP, Kaur A, Singh N. Phenolic compounds as beneficial phytochemicals in pomegranate (*Punica granatum* L.) peel: a review. Food Chemistry. 30 September 2018; 261: 75–86. doi.org/10.1016/j.foodchem.2018.04.039.

55. Bednarczuk VO, Verdam MCS, Miguel MD, Miguel OG. [*In vitro* and in vivo tests used in the toxicological screening of natural products]. Visão Acadêmica [preprint]. 2010 [cited 2019 january 23]. Available from: https://revistas.ufpr.br/academica/article/view/21366/14087 Doi:http://dx.doi.org/10.5380/acd.v11i2.21366. Português.

56. Nguta JM, Mbaria JM, Gakuya DW, Gathumbi PK, Kabasa JD, Kiama SG. Biological screening of kenya medicinal plants using *Artemia salina* L. (Artemiidae). Pharmacologyonline. [preprint] 2011 [cited 2019 january 23] Avalaible from: http://erepository.uonbi.ac.ke/handle/11295/13906.

57. Marreiro RO, Bandeira MFCL, Almeida MC, Coelho CN, Venâncio GN, Conde NCO. [Evaluation of the cytotoxicity of a buccal mouthwash containing Libidibia iron extract] Braz Res Pediatric Dent Int Clinic. [preprint]. 2014 [cited 2019 january 23] Available from: file:///C:/Users/usuario/Downloads/AvaliaodacitotoxicidadedeumenxaguatriobucalcontendoextratodeL.ferrea%20(1).pdf Doi: http://dx.doi.org/10.4034/PBOCI.2014.14s3.04. Português.

